# Spheroid architecture strongly induces miR-221/222 expression and promotes oxidative phosphorylation and survival of mobile tumor cells through a mechanism that includes restriction of miR-9 expression

**DOI:** 10.1101/2023.08.22.554379

**Authors:** Avery S. Ward, Cody N. Hall, Maya O. Tree, D. Stave Kohtz

**Author notes:** Department of Pediatric and Adolescent Medicine, Mayo Clinic, 200 First Street SW, Rochester, MN 55901.

## Abstract

Tumor cell spheroids are three dimensional multicellular structures that form during the expansive growth of carcinoma cells. Spheroids support tumor metastasis as vehicles of dissemination, promoting growth and survival of bulk tumor and cancer stem cells within the mobile tumor cell population. Deciphering how spheroid architecture affects tumor cell phenotype will be essential for the development of therapeutics to inhibit transperitoneal metastasis and the development of peritoneal carcinomatosis by ovarian cancers. We investigated how spheroid formation directly affects OXPHOS activity and microRNA expression in a cultured ovarian carcinoma cell system. The rate of oxidative phosphorylation/respiration per cell in spheroids was nearly double that of the same cell type growing in suspension as single cells. Cells growing as spheroids showed greatly enhanced expression of miR-221/222, an oncomiR that targets multiple tumor suppressor genes, promotes invasion, as well as reduced expression of miR-9, which targets mitochondrial tRNA-modification enzymes and inhibits OXPHOS. Consistent with the greater efficiency of ATP generation afforded by OXPHOS phosphorylation, tumor cells growing as spheroids injected into the nutrient-poor environment of the murine peritoneum survived longer than the cells growing in suspension as loosely associated aggregates. The data suggest that in addition to the reported effects of spheroid formation on cancer cell growth and phenotype, including promotion of stem cell generation, spheroid architecture increases the OXPHOS activity of constituent tumor cells. During the mobile phase of metastasis, when ovarian tumor cells disperse through nutrient-poor environments such as the peritoneum, enhanced OXPHOS activity afforded by spheroid architecture would enhance survival and thereby contribute to metastatic potential.

## Background

The role of three-dimensional architecture in tumor cell growth, invasion, metastasis, and response to therapeutics has been studied using explants derived from tumors, co-cultured mixtures of tumor and normal cells, and tumor cell lines grown in suspension. When directly excised from tumor tissue and grown as mixtures of neoplastic and normal cells, explants are referred to as tumor organoids or tissue-derived tumor spheres [1]. When cultured over nonadherent substrata, some tumor-derived cell lines generate stable spherical or semi-spherical structures referred to as tumor spheroids or tumorspheres [1]. In contrast to loosely associated cell aggregates, which appear as irregular structures, accrue from weak interactions between charged cell surface moieties, and are readily dispersed, tumor cell spheroids are stabilized through homotypic interactions of specific cell surface adhesion proteins [2, 3]. The impact of spheroid formation on tumor cell structure is evident in the smooth spherical morphology, polarity, and contractile behavior of spheroids [4]. The effects of the spheroid architectural on gene expression and phenotype of constituent tumor cells are evident in enhanced development of cancer-initiating or cancer stem cells (CSCs) [5] [6], enhancement of drug-resistance and survival [7, 8], and increased tumorigenicity and invasion in animal models [4]. The significance of tumor cell spheroid formation in cancer progression and metastasis has emerged from several studies [9]. Tumor cell spheroids are cognate multicellular vehicles that mediate some forms of metastasis. Spheroid formation by cultured ovarian cancer cells has been shown to promote invasive behavior in vitro and in vivo [4, 7, 10], indicating that tumors that spread by traversing intraperitoneal spaces likely exploit spheroid formation to facilitate mobility and survival while metastasizing to and invading distal sites (https://pubmed.ncbi.nlm.nih.gov/19132753/). Epithelial ovarian cancers frequently metastasize intra-abdominally, often forming numerous smaller sized lesions that are virtually impossible to remove during surgical debulking. This condition is referred to peritoneal carcinomatosis, and is associated with a high rate of post-treatment recurrence [11].

The formation of tumor spheroids in vivo is thought to proceed either from collective detachment of groups of cells from the primary tumor, or by growth of spheroids from individual or small groups of cells shed by the tumor [10]. The sequence of events leading to spheroid formation in vitro varies between different tumor cell lines [12]. Some tumor cell lines, after being grown as adherent monolayers on tissue culture plastic and dissociated enzymatically into single cells, spontaneously associate to form coherent spheroids when cultured over ultra-low attachment surfaces. For other cell lines, proliferation of single or small number of dissociated cells results in the formation and progressive enlargement of spheroidal structures. The changes in cell architecture, polarity, and contractility, and the changes in gene expression and cellular phenotype associated with spheroid formation are directly or indirectly driven by the homotypic interactions of cadherins and the formation of adherens junctions [3]. Homotypic binding also modulates interactions of the intracellular cytoplasmic tails with actin filaments and with adaptors that mediate intracellular signaling [13]. Differential expression of E, N, and/or K-cadherin genes (*CDH1*, *CDH2*, *CDH6*) is directly involved in stabilizing cell-to-cell interactions required for spheroid formation [14]. Spheroid formation can be blocked by exposing tumor cells to inhibitory antibodies or small molecules that target expressed cadherin isoform(s) [14]. A comparison of epithelial-mesenchymal transition (EMT) in different cancer cell lines revealed a spectrum of phenotypes, with ovarian cell lines aligning into an intermediate group that preferentially expressed N-cadherin over E-cadherin and was the most spheroidogenic and anoikis-resistant of the tested groups [15].

In traversing the peritoneum or internal spaces, mobile epithelial ovarian tumor cells are challenged by lack of substrate for adhesion and by restricted access to nutrients [16]. We considered how spheroid formation allows mobile tumor cells to overcome these challenges; in particular, how spheroid architecture affects the efficiency of oxidative phosphorylation (OXPHOS). It is difficult to answer this specific question by comparing different tumor cell lines with different innate spheroid-forming properties as differences in OXPHOS will be conferred by genetic and epigenetic variations inherent to cell lines derived from different tumors. While studies of this type have indicated that enhanced OXPHOS promotes greater or more stable spheroid formation [17], it is not clear from these studies how spheroid formation impacts the OXPHOS activity of tumor cells. In addition, comparisons of spheroidal and dissociated forms of a single tumor cell line are distorted by effects of trypsin or other non-specific mediators of dissociation on OXPHOS activity [18, 19]. To address these issues, we derived subcultures from the high-grade endometrioid or dedifferentiated epithelial ovarian tumor cell line TOV-112D [20] that grow over ultralow attachment polystyrene either as spheroids or as dispersible clumps of single cells. The TOV-112D cell line does not express E-cadherin (*CDH1*; [21]) and evidence is presented that spheroid-forming activity of TOV-112D cells is modulated by differential expression of N-cadherin. Spheroid formation by TOV-112D cells growing over ultralow attachment plastic (ULP) was found to enhance OXPHOS activity and strongly increase expression of miR-221/222 compared TOV-112D growing as loosely associated clumps of cells. Enhanced OXPHOS in spheroids was found to be due in part to repression of miR-9-3p. In contrast to the view that the Warburg Effect reflects a universal reliance of tumor cells glycolytic metabolism, recent studies have indicated that tumor cells in nutritionally deficient microenvironments are acutely dependent on mitochondrial function [22]. Consistent with a role for mitochondrial function in the survival of mobile tumor cells, *in vivo* experiments presented here showed that spheroid forms of TOV-112D cells display greater capacity for intraperitoneal survival than non-spheroidal cells.

## Materials and methods

### Materials

Early passage TOV-112D cell line was obtained from the ATCC. Janus Green B was obtained from Millipore Sigma (210677). Antibodies: Anti-rabbit fluorescein (Vector Labs, FI-1000-1.5); Alpha-tubulin (Cell Signaling, 2144); Citrate synthetase (Abcam, ab96600); Cytochrome c (Abcam, ab13575); GAPDH (Novus Biologicals, NB100-56875); GTPBP3 (Proteintech, 10764-1); MTO-1 (Proteintech, 15650-1-AP); N-Cadherin (Invitrogen, 33-3900); OXPHOS panel (Abcam, ab110413); p38 (Santa Cruz); TRMU (Invitrogen, PA5-57208) TRMT1 (Santa Cruz, SC-373687).

### Culturing and counting of cells

TOV-112D and derivatives were cultured in DMEM supplemented with 15% fetal bovine serum, high glucose (4.5 g/L), 1 mM pyruvate, 3.9 mM L-alanyl-glutamine, 100 uM non-essential amino acids, and 20 ug/ml gentamicin. Ultralow attachment culture vessels were purchased from Corning (Six well plates, part number 3471; 24 well plates, part number 3473). The growth rates of TOV-LT and CAP cells grown in long-term pass cultures over ULP were indistinguishable. For a typical experiment, TOV-LT cells were cultured over ULP, counted using a Neubauer counting chamber (.1 mm depth), and distributed into MiR05 buffer at 0.5 – 1.0 × 10^6^ cells per ml. CAP cells were grown on TCP, dissociated with trypsin and replated at 1:20 of final density over ULP. After freshly dissociated CAP cells were grown for 4-5 days small spheroids (16-32 cells) that were counted using the Neubauer chamber. Cultured that contained larger spheroids were counted using a Malassez counting chamber (.2 mm depth). All cell counts were performed before and after Oxygraph-2k respirometry, and the MiR05 combined with digitonin and mechanical stirring performed during the assays rendered the cultures into single cells.

### Phase and immunofluorescence microscopy

Phase contrast microscopy of live cells cultured on tissue culture plates was performed using a Nikon Diaphot microscope. For immunofluorescence microscopy, cells cultured suspension over ULP were, pelleted by centrifugation, resuspended in PBS, fixed with with equal volume fresh 4% paraformaldehyde in PBS. Paraformaldehyde was quenched with 50 mM glycine in HMK buffer (20 mM HEPES, pH 7.5; 1 mM MgCl_2_; and 100 mmol/L KCl). After cells were treated with 0.5% Triton X-100 in HMK, they were incubated with blocking solution (10% normal donkey serum and 1% bovine serum albumin in HMK buffer). Cells were incubated with primary antibody for 1 to 2 hours at room temperature or overnight at 4°C. After washing with HMK five times, the cells were incubated with fluorescent secondary antibodies. Cells were washed in PBS and mounted with VectaShield mounting medium with DAPI (Vector Laboratories, Burlingame, CA). Fluorescence microscopy was performed using a Leica DM5000 microscope.

### Lipofection

Cells were plated during log phase growth in fresh medium over ULP for 24 hours, then transfected with small-interfering RNA (siRNA) using Lipofectamine RNAiMAX (Invitrogen, Carlsbad, CA) or Lullaby (OZ Biosciences, San Diego, CA) and the manufacturer’s recommended procedures. Assays were performed 48-72 hours after lipofection. Sources of siRNA: Control siRNA A or B (Santa Cruz Biotechnology, (SC-37007 or SC-44230); CDH2 (Santa Cruz Biotechnology, SC-29403); MTO1 (Santa Cruz Biotechnology. Anti-miRs were lipofected with RNAiMAX or Lullaby. Anti-miRs 9-5p and 9-3p were purchased from Accegen; anti-miR 9-5p and 9-3p were combined for experiments designated anti-miR-9.

### Differential expression analyses and quantitative PCR

Libraries were prepared from enriched small RNA (smRNA) isolated from TOV-LT and CAP cells grown over ULP. Libraries were sequenced on a Hi-Seq X10 instrument and normalized to 1 million total reads by Omega Bioservices, Norcross GA (Supplementary Figure 2). Secondary screening is ongoing, with samples showing either 10-fold difference in signal or on/off expression tested further for significance by quantitative PCR.

Total RNA was isolated from cells using the miRNeasy minikit (Qiagen). Quantitative PCR was performed using the Mir-X miRNA qRT-PCR TB Green Kit (Takara Bio, USA) using the protocol provided by the manufacturer. The cDNA was isolated with a kit from Zymo Research (D7011), quantified by nanodrop spectrophotometry, and 5 ng was used per qPCR reaction. Quantitative PCR was performed using the Applied BioSystems StepOnePlus real-time PCR system. Because equal amounts of cDNA generated from sample RNAs were amplified, both delta-Ct (dCt) and delta-delta (ddCt) values using U6 RNA as control were reported. Relative expression calculations: When × has the lower cycle threshold (Ct), relative expression was determined as [miR-X]/[miR-Y] = 2^Ct(miR-Y)^/2^Ct(miR-X)^ ; to use U6 as normalization standard: [U6-X]/[U7-Y] = 2^Ct(U6-Y)^/2^Ct(U6-X)^ = normalization factor. Multiplication of miR relative expression by the U6 normalization factor produced the same outcomes as methods that normalizes the Ct values of the miR with those of U6 to give dCt values. At least three independent qPCR determinations were performed from a cDNA preparation, and the experimental comparisons were repeated from independent cultures of cells. The mRQ 3’ primer was used as the reverse primer for the miR qPCR reactions.

**Table 1.**
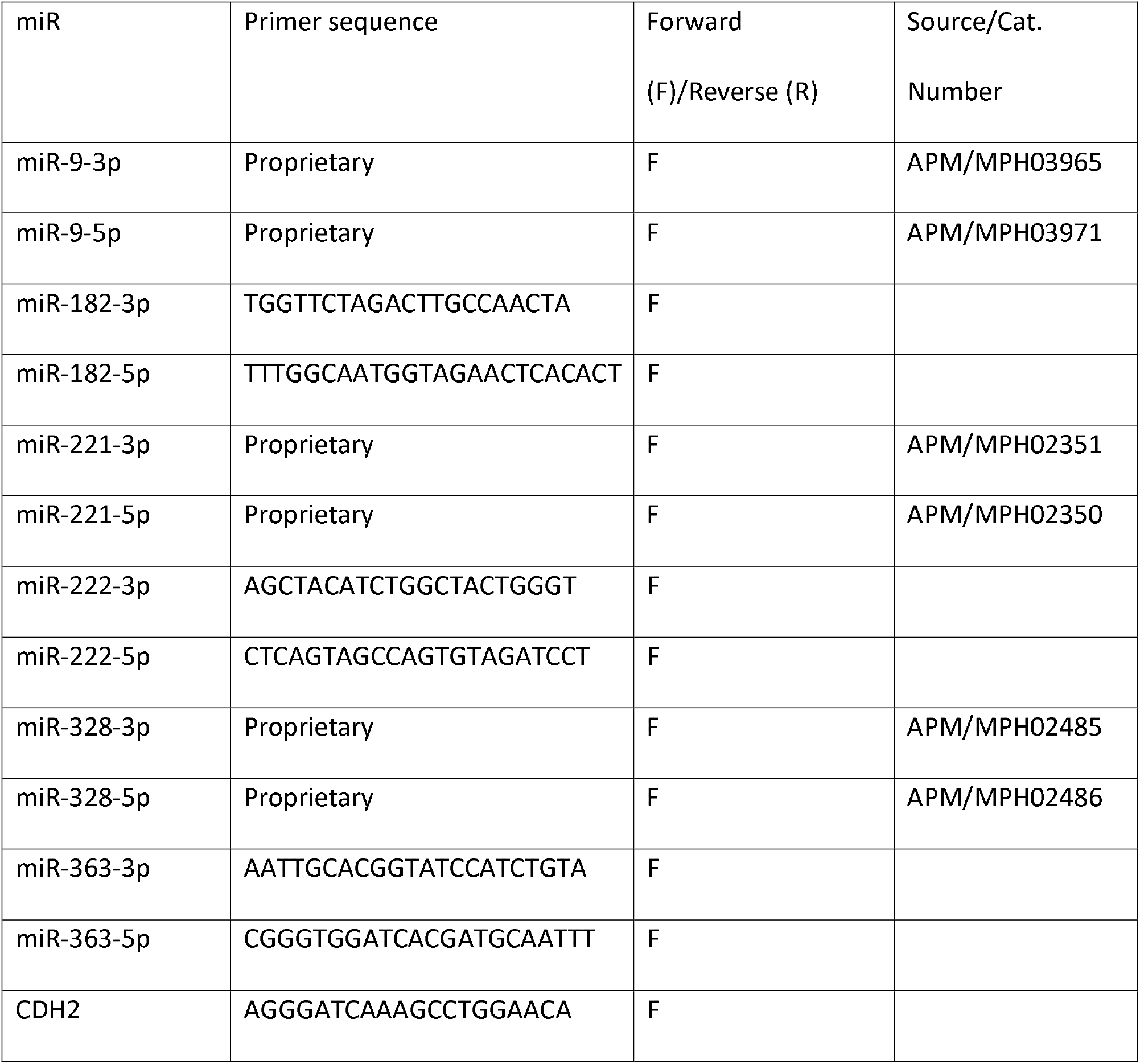
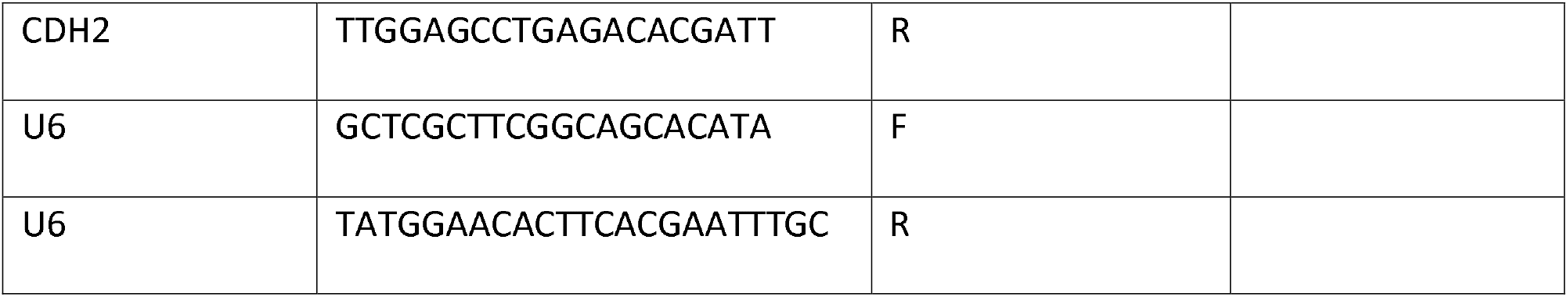
Sequences of primers used to detect miR, CDH2 mRNA, and U6 RNA in qRT-PCR analyses.

### Western blot analyses

Whole cell lysates were prepared by washing cells once with PBS, followed by direct suspension of plates in SDS sample buffer (50 mmol/L Tris-HCl, pH 6.8; 10% glycerol; 2% SDS; 5% β-mercaptoethanol; 125 μg/mL bromophenol blue). Samples were removed, incubated at 80° C, and flash frozen in liquid N_2_. Samples were flash thawed at 80° C immediately before loading. Sodium-dodecyl-sulfate polyacrylamide gel electrophoresis and transfers of proteins to .45uM nitrocellulose membranes were performed using the Protean 2 or 3 systems (Bio-Rad) and protocols provided by the manufacturer. Blots were blocked, incubated with primary antibody, and incubated with fluorescent secondary antibodies as described for quantitative western blots by Li-Cor Biosciences. Western blots were visualized and quantified using a Li-Cor Odyssey Fc imager and software. Experimental determinations were repeated three times in parallel lanes on the same blot, and compared for significance by Student’s T-test.

### Respirometry

Equal numbers of cells (1-2 × 10^6^) were suspended in 2 ml of mitochondrial respiration medium (MiR05: 110 mM sucrose, 60 mM K+-lactobionate, 0.5 mM EGTA, 3 mM MgCl_2_, 20 mM taurine, 10 mM KH_2_PO_4_, 20 mM HEPES adjusted to pH 7.1 with KOH at 37 °C; and 1 g/l BSA). Cells were permeated by the addition of digitonin (10 ug/ml). A two-channel titration injection respirometer (Oxygraph-2k, Oroboros Instruments) was used to analyze liver respiratory function after homogenization (temperature was 28°C). Oxygen flux (pmol/(s*ml)) and oxygen concentration (μM) were recorded in real-time using Oroboros Datlab software (Oroboros Instruments). Routine respiration experiments were conducted as shown in Figure 3. After addition of malate (2 mM) and pyruvate (5 mM), the OXPHOS capacity of complex I (CI) was examined after adding ADP (5 mM). Succinate (10 mM) was added to measure OXPHOS of CI and complex II (CI&II). Addition of carbonyl cyanide 4-trifluoromethoxy)phenylhydrazone (FCCP, 2 μM) was performed at least twice to obtain the maximal uncoupled respiratory capacity of the electron transfer system (Uncoup). Finally, antimycin A (1 μM) was added to evaluate residual oxygen consumption (which was not detected in our experiments). Oxygen flux values were compared for equal cell numbers (1 × 10^6^ or 2 × 10^6^ cells as indicated) per two ml assay volume. Pair of determinations on the Oxygraph-2k were repeated with independent sets of cells at least three times, and normalized by cell number. Data were analyzed for standard error and for significance by two-way unpaired ANOVA.

### In vivo survival experiments

Experimental testing of TOV-LT (dispersible clumps) and CAP (spheroid) intraperitoneal (i.p.) survival in mice was performed by Altogen Labs (Austin, TX) under IACUC study protocol LC03516. NOD/SCID (NU(NCr)-Foxn1nu mice (10-13 weeks) were purchased from Harlan Laboratories. To confer GFP expression, CAP or TOV-LT cells for transduced with defective lentivirus (EF1a-Luciferase-2A-GFP;) and selected with bsd. Transduced CAP or TOV-LT cells for injection were grown for 4 days over ULP, diluted into new medium over ULP and then harvested in log growth phase. At this stage spheroid growth is readily apparent in CAP cultures and absent from TOV-LT cultures. Acclimated mice were each injected i.p. with a single dose of 10 million cells (six mice with CAP cells and six mice with TOV-LT). In vivo imaging was performed with the Night Owl imaging instrument with associated Indigo software (Berthold Technologies). Luciferase expression was analyzed after intraperitoneal luciferin injection as described in Berthold Technologies application notes. 125 mg/kg (bodyweight) of D-Luciferin was administered i.p. 10 minutes prior to making luciferase measurements. Animals received Ketamine/Xylazine anesthesia. Two types of signals were recorded - photo and luminescence. A positive control of 0.5 U87-Luc cells was injected subcutaneously into a non-experimental animal was used to validate the luminescence reaction, and an uninjected mouse was used to determine the background threshold. Mice were imaged every 5 days after injection for 40 days. Six mice were injected with TOV-LT cells and six mice with CAP cells. No animals were lost during the course of the experiment. Significance was determined was determined after comparing signal intensities by two-way ANOVA.

## Results

### Isolation and characterization of subcultures of a clonal tumor cell line that grow in suspension as individual cells or as spheroids

The TOV-112D tumor cell line was originally derived from a patient with in a histological diagnosis of endometrioid ovarian carcinoma [23], although this designation was later revised to be both endometrioid and dedifferentiated [20]. Genetic analyses showed that TOV-112D cells contains an activating mutation in CTNNB1 (β-catenin) a common defect in low-grade endometrioid ovarian carcinomas, as well as mutated TP53, a characteristic of high-grade ovarian carcinomas. In contrast to the more common early TP53 defects that drive development of high grade ovarian serous adenocarcinomas, defects in the TP53 alleles of TOV-112D cells were likely acquired late in tumor development, possible as a clonal outgrowth from benign or low-grade tumor cells [24].

Vials of frozen TOV-112D cells were originally obtained from ATCC and stocks were generated from adherent cells grown and expanded on tissue culture plastic. TOV-112D cells grow as epithelial-like cells on tissue culture polystyrene (TCP, Fig. 1A). When dissociated from tissue culture polystyrene by trypsin and passed to ultra-low attachment polystyrene (ULP), early passage TOV-112D cells grow as a mixture of spheroid and non-spheroidal cells. Rather than forming spontaneously from aggregates of cells, nascent spheroids gradually grew out from single or small groups of cells over a few days from TOV-112D cells cultured over ULP. On day 5 of culture over ULP, small refractile groups of cells (spheroids) are apparent (Fig. 1B, arrows). After passage, on day 9 of culture over ULP, the nascent groups of refractile cell have grown into larger spheroids with smooth reflective exterior and a darker interior visible under phase microscopy (Figs. 1C and 1D, arrows). When grown continuously over ULP, TOV-112D cells gradually loose spheroid-forming capacity, and after five or more passages grow only as individual cells in loosely associated aggregates (Figs. 1E and 1F). Cultures of TOV-112D grown as individual non-spheriodal cells in this manner are referred to as TOV-LT cells. A clone of TOV-112D cells that had been modified in other studies to express increased nucleoporin p62 (NUP62) in response to treatment with doxycycline has been reported earlier as TOV-112D-CAP cells [25]. In the absence of doxycycline, TOV-112D-CAP (referred to herein as CAP) cells express NUP62 at levels equivalent to parent TOV-112D cells. We found that CAP cells display stable growth over ULP predominantly as spheroids for multiple (10+) passages (Figs. 1G and 1H). Cultures of TOV-LT (loose aggregates) and CAP (spheroids) cells grown in parallel over ULP were passed at the same intervals and at equivalent dilutions for several generations, suggesting that their rates of growth were similar.

**Figure 1.**
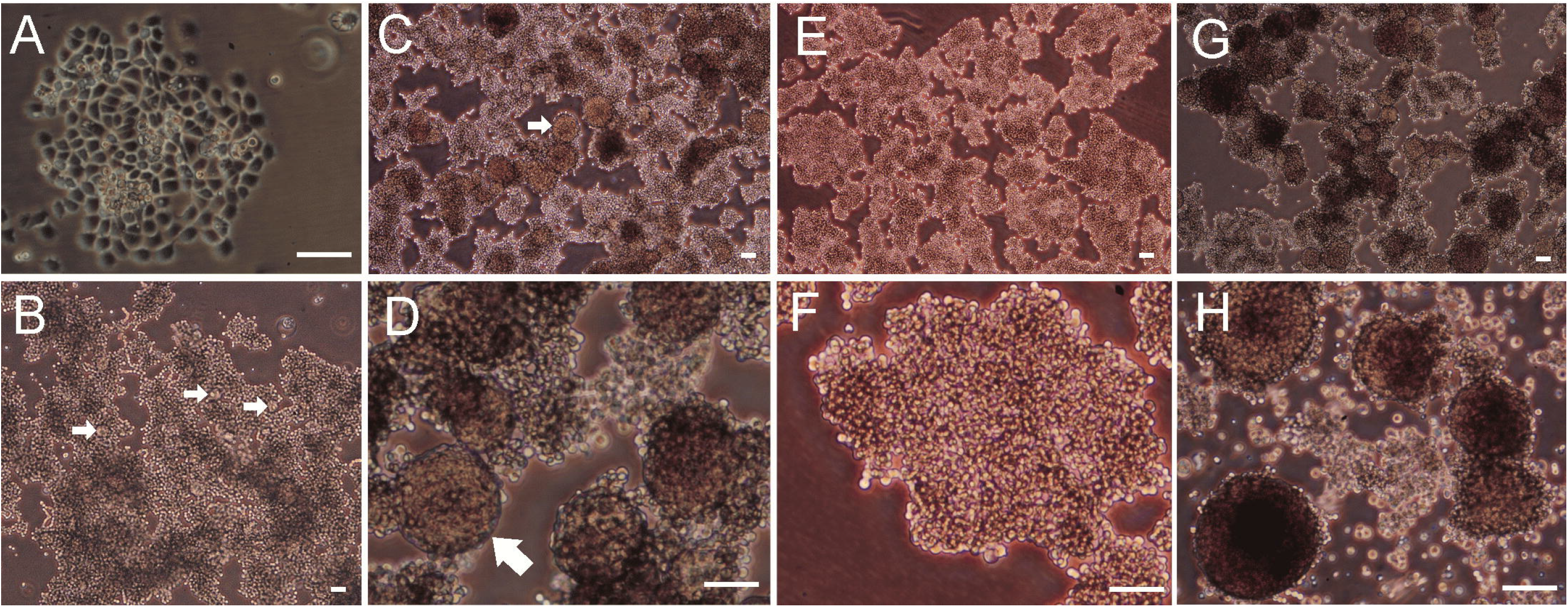
Phase microscopy of parent TOV-112D cells, TOV-LT subcultures, and CAP cells in culture. A. TOV-112D cell grown as adherent cells on tissue culture polystyrene (TCP). B. TOV-112D cells 4 days after first passage over ultralow attachment polystyrene (ULP, day 5); nascent spheroids are indicated by arrows. C. TOV-112D cells 4 days into second pass over ULP (day 9); growth of large spheroids is observed (arrow). D. Higher magnification of spheroids shown in C (arrow); E. TOV-112D cells after 5+ passages over ULP; these cultures are referred to TOV-LT cells. Spheroids are absent and cells are growing in loosely associated aggregates. F. Higher magnification of cells in E showing absence of spheroids. G. CAP cells grown for several passages over ultralow attachment plastic showing abundant growth of large spheroids. H. Higher magnification of spheroids from culture shown in G. Scale Bar indicates 50 um.

### Expression of N-cadherin is associated with spheroid formation by TOV-112D cells

Early passage TOV-112D cell cultures were grown on TCP, dissociated with trypsin, and cultured over ULP. The cells were passed continuously over ULP under two media conditions (R - RPMI or D - DMEM) and sampled at progressively later passage numbers for analysis by Western blot with an N-cadherin antibody. Early passage TOV-112D cultures grown on TCP in either medium showed accumulation of N-cadherin (Fig. 2A, adh. lanes), but N-cadherin (*CDH2*) expression was lost after six passages (Fig. 1A, sus. p6). The loss of N-cadherin expression at passage six (p6) corresponds to the loss of spheroid formation (Fig. 1), with TOV-112D cultures grown over ULP at this passage or beyond referred to as TOV-LT cells. CAP cultures continue to express N-cadherin (Fig. 2A, p6+) and continue to show spheroid formation at passage six and beyond (Fig. 1G). Comparison of N-cadherin transcript levels in TOV-LT and CAP cells by qPCR showed 16.5 times greater expression of the *CDH2* gene in CAP cells (Fig. 2B).

**Figure 2.**
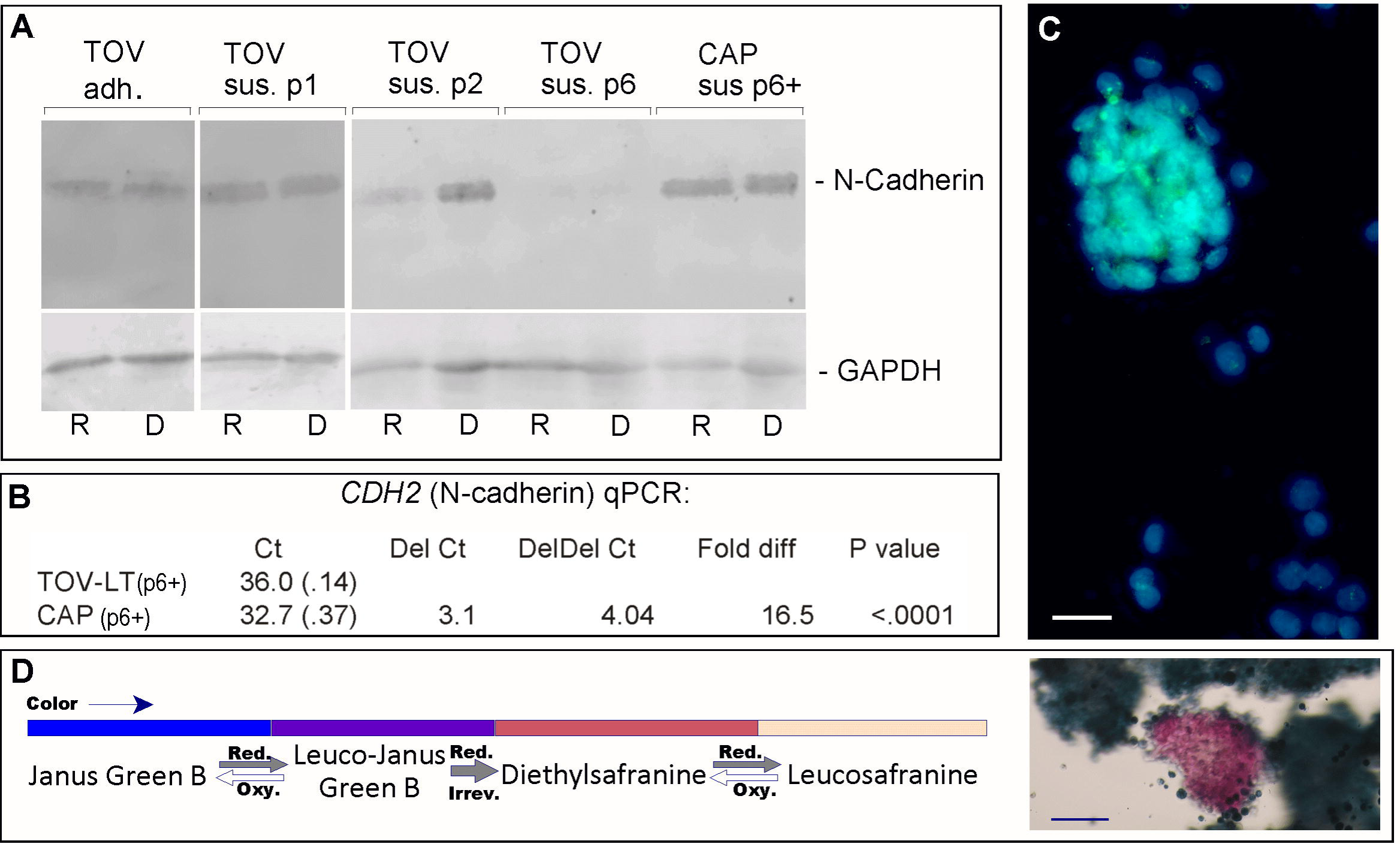
Expression of N-cadherin (*CDH2*) is directly associated with spheroid-forming ability of TOV-112D cells. A. Western blot analysis of N-cadherin expression in TOV-112D (TOV) cells grown as adherent (adh.) cultures on TCP and as suspension (sus.) cultures over ULP. Two media type were used (R - RPMI and D - DMEM), and passage numbers are indicated (e.g., p1). CAP cells (described in text) are clonal variants of TOV-112D that retain spheroid-forming ability after several passages. B. Quantitative PCR comparison of *CDH2* gene expression in TOV-LT and CAP cells cultured over ULP. C. N-cadherin immunofluorescence (green) of CAP cells cultured over ULP. Nuclei are stained with Hoechst 33342 dye. Bar indicates 25 um. D. Left: Diagram indicates color changes associated with reduction of Janus Green B, Leuco-Janus Green B, or Diethylsafranine. Right: Janus Green B stain of spheroid and clumped CAP cells growing over ULP. Bar indicates 100 um.

Immunofluorescence microscopy of CAP cells grown over ULP with N-cadherin antibody revealed a course granular staining pattern that was most intense in the spheroids (Fig. 2C). Individual cells growing outside of spheroids showed lower levels or undetected immunofluorescence. CAP cells were grown over ULP in the presence phenazine dye Janus Green B (JG-B) which, as a vital stain, can be used to show relative mitochondrial function [26]. Reduction of blue-green JG-B by mitochondrial dehydrogenase activity results in the production of diethyl safranine, which is bright pink. Conversion of JG-B to diethyl safranine first involves reversible generation of the intermediate Leuco-Janus Green B; reduction of Leuco-Janus Green B to diethyl safranine is essentially irreversible (Fig. 2D). Cells with greater mitochondrial dehydrogenase activity will more rapidly accumulate diethyl safranine, and within a specific window of time after treatment will appear pink in comparison to the blue-green color of slower respiring cells. Among CAP cells cultured over ULP, some large spheroids structures stained pink/red after a 15-minute incubation with JG-B at 37° (Fig. 2D). In contrast, clumps of TOV-LT cells or of CAP cells that did not form spheroids stained blue after incubation with JG-B under the same conditions. This observation suggests that tumor cell growth as spheroids is associated with increased mitochondrial OXPHOS activity compared to cells growing as loosely associated clumps.

### Formation of spheroids enhances OXPHOS activity of tumor cells growing in suspension

The rates of OXPHOS of cells growing over ULP as loose aggregates (TOV-LT) or as spheroids (CAP) were compared by high resolution respirometry on digitonin-permeated cells. The growth rates of CAP and TOV-LT cells over ULP were found to be similar several passages, but later passages CAP spheroids become large and difficult for cell counts. For respirometry experiments, CAP cells were cultured as monolayers over TCP, then digested with trypsin, counted and cultured in parallel with TOV-LT cells over ULP at the same seeding density. After 5 days of culture, TOV-LT and CAP cells grew to similar densities, but small to moderate-sized spheroids were formed only in the CAP cultures. After counting, these TOV-LT and CAP cells were used for respirometry experiments. For a typical experiment, equal cell counts (1-2 × 10^6^ cells per 2 ml) of TOV-LT (loose aggregates) or CAP (spheroids) cells were suspended in MiR-05 buffer [27], treated with digitonin, and incubated with pyruvate and malate. Complex I (CI)-linked state 3 respiration was initiated by the addition of ADP; Complex I + II (CI+II)-linked state 3 respiration was initiated by the addition of succinate; and uncoupled respiration was initiated by the addition of FCCP (Figs. 3A and B). Signal specificity from OXPHOS was confirmed by inhibition to baseline after addition of antimycin A. Activities were quantified from the peak rates of O_2_ flux after each treatment, and the mean values from five independent experimental determinations were compared (Fig. 3C). CAP cultures showed significantly greater CI, CI+CII, and uncoupled state 3 (3U) respiration rates per cell than TOV-LT cultures. This result suggests that growth in spheroid form enhances OXPHOS activity in comparison to growth in suspension as individual or dispersible clumps of cells. Western blot analyses of a panel of proteins representing four of the mitochondrial respiratory chain complexes did not reveal overt differences between TOV-LT and CAP cells (Fig. 3D). Comparison of citrate synthase, cytochrome c, and GAPDH revealed increased accumulation of cytochrome c in CAP cells (Fig. 3D).

**Figure 3.**
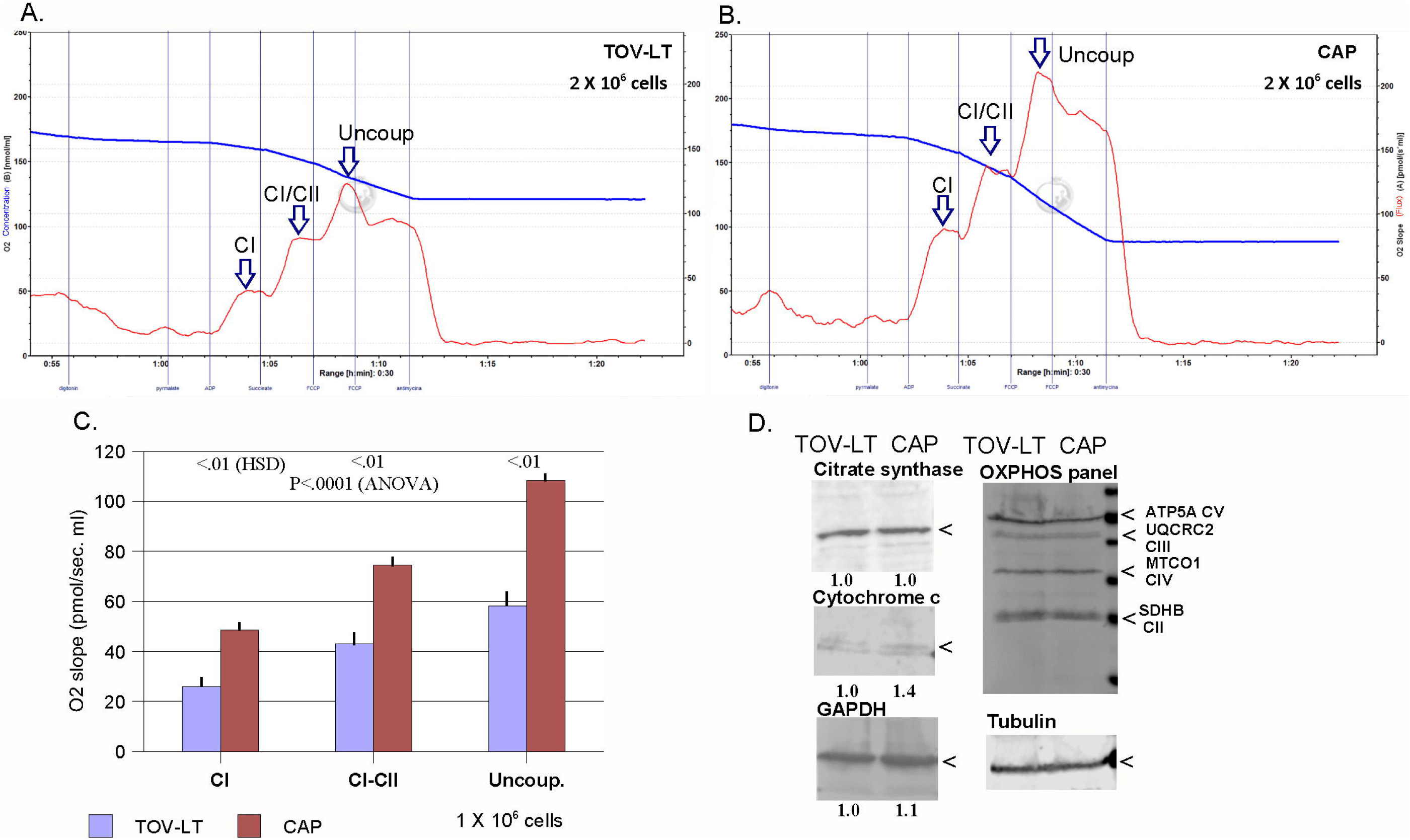
Comparison of OXPHOS activity in TOV-LT and CAP cells growing over ULP. A and B. Example of comparative respirometry plots of 2X10^6^ TOV-LT (A) or CAP (B) cells. Samples were run in parallel on the Oroboros O2k respirometer. Blue plot shows total O_2_ consumption; red plot shows O_2_ flux (continuous change). Time points of addition of digitonin, pyruvate and malate (pyrmalate), ADP, succinate, FCCP, and antimycin A. Positions of flux data points for CI, CI+CII (CI/CII) and uncoupled respiration are indicated by arrows. C. Mean values of five independent respirometry determinations for TOV-LT and CAP cells. Results of ANOVA and Tukey HSD tests are shown. D. Western blot analysis of whole cell lysates from equal numbers TOV-LT and CAP cells grown over ULP. Relative quantification of citrate synthase, cytochrome c, and GAPDH normalized to tubulin in TOV-LT and CAP cell lysates is shown.

We sought to determine how experimental dissociation of CAP spheroids affects OXPHOS rate. When CAP or TOV-LT cells were cultured on TCP, released by trypsin/EDTA digestion, and assayed immediately thereafter by respirometry, OXPHOS activity was greatly suppressed in both cell suspensions (Supplementary Fig. 1). In contrast to results reported for human dermal fibroblasts [28], trypsin/EDTA digestion would be not a useful technique for detachment of TOV-112D carcinoma cells, with comparisons of respiration from intact and trypsin/EDTA-digested spheroids most likely to yield artifactual differences. As shown above, early passage TOV-112D cells and CAP cells persistently express N-cadherin that is directly associated with spheroid formation over ULP, and lack expression of E-cadherin (*CDH1*). We next determined how treatment of CAP cells with siRNA directed at CDH2 transcripts affects the integrity of spheroid structures. CAP cells cultured in DMEM or RPMI over ULP were lipofected with siRNA directed at CDH2 (N-cadherin) transcripts, and western blot analysis showed overt reduction of N-cadherin levels in comparison to cells cultured with control siRNA (Fig. 4A). Examination of transfected cultures by phase microscopy showed dissolution of spheroids, with remnants often appearing in crudely formed clumps (Fig. 4B). Visible quantification of spheroid formation showed that cultures transfected with CDH2 siRNA contained about 60% less spheroids than were present in cultures treated with control siRNA (Figure 4C). Treatment with CDH2 siRNA also disrupted the structure of remaining spheroids in a manner that is not readily apparent from visual inspection. We next determined how inhibition of spheroid formation by siRNA directed at CDH2 transcripts affects OXPHOS activity of CAP cell spheroids. Respirometry of CAP cells 72 hours after lipofection with CDH2 siRNA showed significant reduction of Complex I, Complex I/II, uncoupled state 3 respiration (Fig. 4D) compared to CAP cells lipofected with control siRNA. These data indicate that the spheroidal architecture, which is disrupted by siRNA directed at CDH2, supports increased OXPHOS activity.

**Figure 4.**
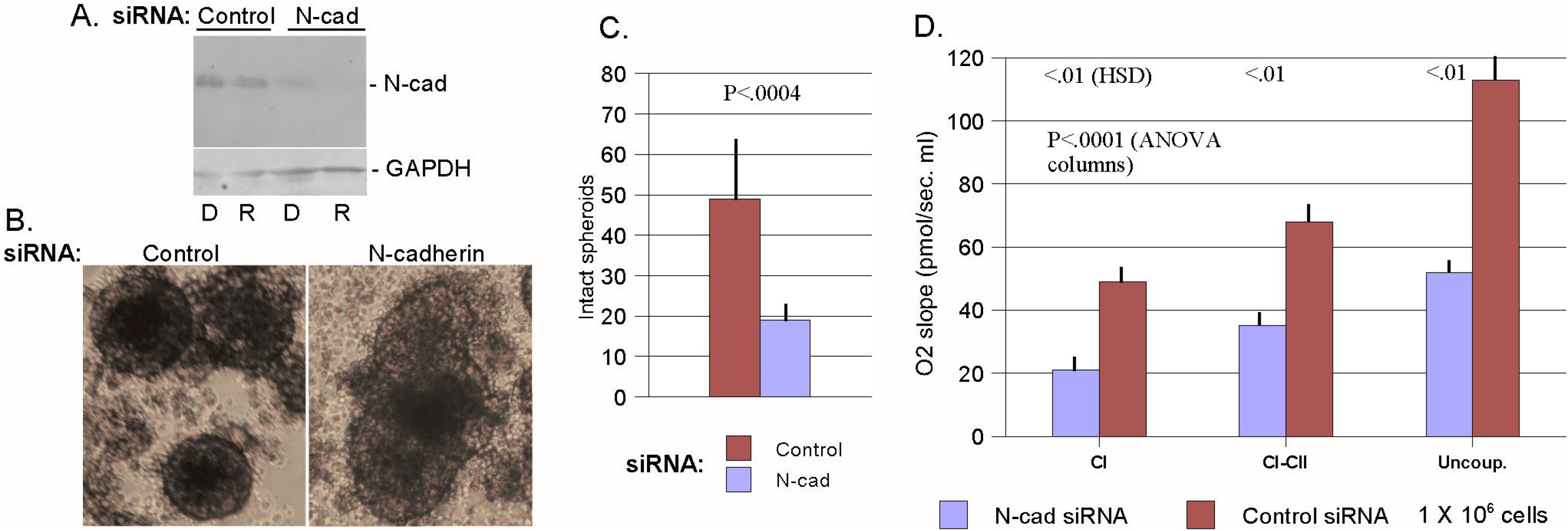
Inhibition of N-cadherin expression disrupts spheroid architecture and reduces OXPHOS activity of CAP cell. A. Western blot analysis of N-cadherin accumulation in CAP cells grown over ULP 72h after lipofection of Control or CDH2 siRNA (N-cad). Western blot analysis of the same blot with an antibody to GAPDH is shown as a loading control. B. Phase microscopy of CAP cells grown over ULP 72 hours after lipofection of Control of CDH2 siRNA (N-cadherin). C. Relative counts of intact spheroids in CAP cultures grown over ULP 72 hours after treatment with Control or CDH2 siRNA (N-cad). The mean number of intact spheroids observed in several low power fields is shown with the standard error and T-test P-value. D. Comparative respirometry of CAP cells grown over ULP 72 hours after lipofection with Control or CDH2 siRNA (N-cad). Mean values of five independent respirometry determinations for CDH2 or control siRNA-treated cells are shown with standard errors. Results of an ANOVA and Tukey HSD test are shown.

### Differential expression of microRNAs in spheroidal and non-spheroidal TOV-112D cells

Libraries were prepared from enriched small RNA (smRNA) isolated from TOV-LT and CAP cells grown over ULP. Libraries were sequenced on a Hi-Seq X10 instrument and normalized to 1 million total reads (Supplementary Fig. 2). From this initial screen, candidate microRNAs (miRs) were identified that were further analyzed for relatively enhanced expression in TOV-LT or CAP cells by quantitative reverse transcriptase PCR (qRT-PCR). MicroRNAs verified as upregulated in CAP spheroids include miR-221/222 (Fig. 5A) and miR-328 (Fig. 5C). Also shown on the same scale is elevated expression of *CDH2* detected in CAP cells. Compared to TOV-LT cells, highly enhanced expression of miR-221/222 was observed in CAP cells and intermediate expression was observed in first passage of TOV-112D cells (TOV-1) over ULP (Fig. 5B). As indicated above, early passage TOV-112D cells maintained on TCP form generate spheroid in early passages over ULP (TOV-1 cells), losing the ability to form spheroids after five passages over ULP (TOV-LT). Expression of miR-221/222 in TOV-1 cells was observed at a level between that of TOV-LT and CAP cells is consistent with a direct association with spheroid formation. Expression of miR-182, miR-9, and miR-363 were observed to be higher in TOV-LT than in CAP cells grown over ULP, suggesting a negative association of expression of these microRNAs with spheroid formation (Fig. 5D).

**Figure 5.**
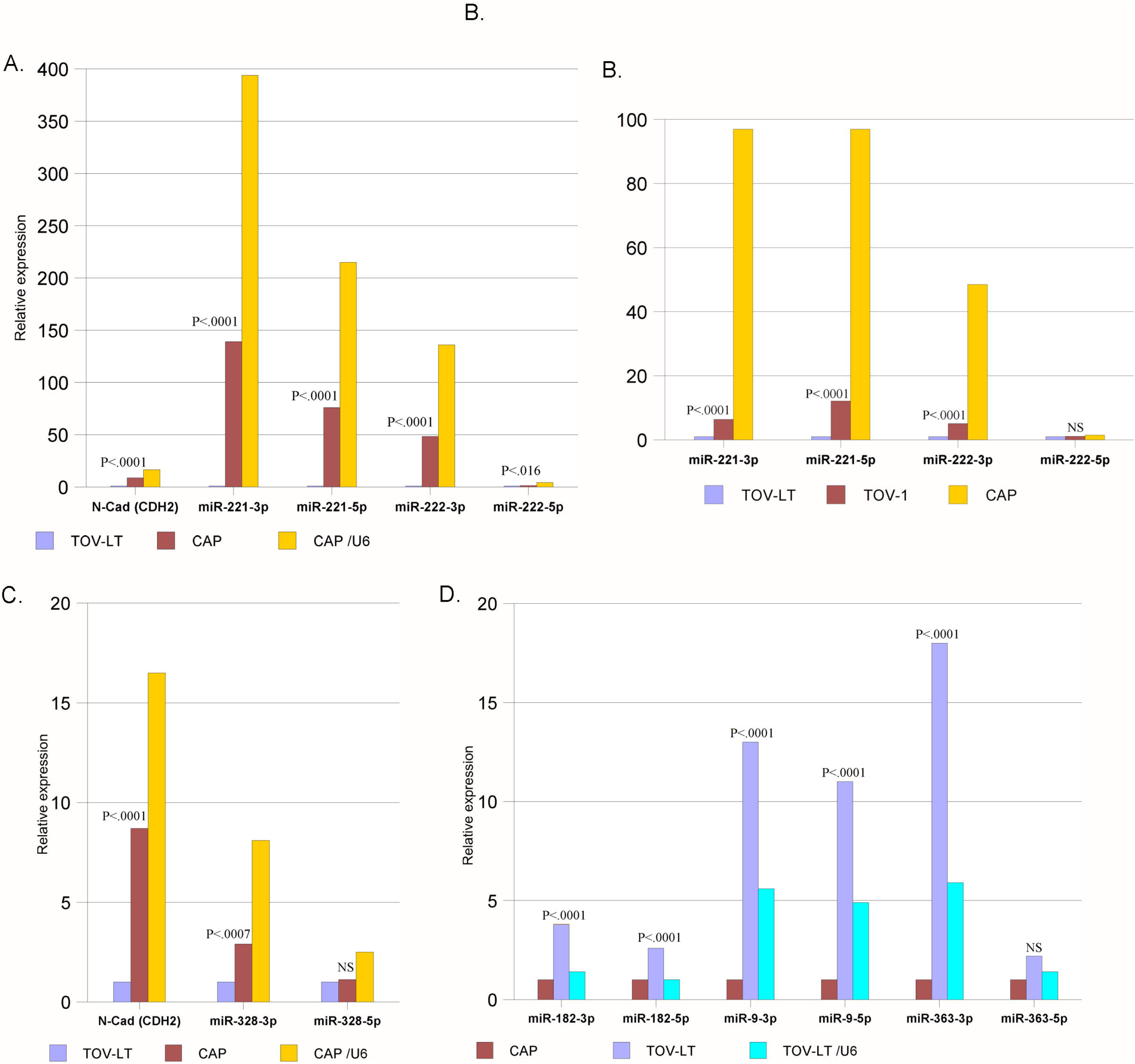
Differential expression of microRNAs in TOV-LT and CAP cells. Quantitative reverse transcriptase PCR (qRT-PCR) analysis was performed on microRNAs isolated from TOV-LT or CAP cells grown over ULP. The quantity of cDNA used in each reaction was 5 ng. Mean results of three or more independent determinations performed in parallel are shown with P values derived from Student’s test (T-test). A. Relative expression levels of N-cadherin (CDH), and miR-221/222 in CAP cells grown over ULP. Values are relative to TOV-LT. B. Relative expression levels of miR-221/222 in early passage TOV-112D (TOV-1), TOV-LT, and CAP cells cultured over ULP. Values are relative to TOV-LT. Relative values normalized to U6 are also shown. C. Relative expression levels of N-cadherin (CDH2), and mR-328 in CAP cells grown over ULP. Values are relative to TOV-LT. D. Relative expression levels of miR-182, miR-9, and miR-363 in TOV-LT cells grown over ULP. Values are relative to CAP cells. Relative values normalized to U6 are also shown, although TOV-LT cells consistently showed higher U6 levels.

Increased expression of miR-9 has been linked to reduced mitochondrial function via down-regulation of MTO1, GTPBP3, and TRMU [29, 30], genes that function in mitochondrial tRNA modification. To consider the effects of enhanced miR-9 expression in TOV-LT cells, respirometry experiments were performed with TOV-LT cells cultured over ULP and lipofected with antagomirs (anti-miRs) to both forms of miR-9 (5p + 3p). Treatment of TOV-LT cells with miR-9 anti-miRs resulted in significant enhancement of Complex I, Complex I/II, uncoupled state 3 respiration compared to TOV-LT cells lipofected with control siRNA (Fig. 6A). Western blot analysis of whole cell lysates of TOV-LT cells grown over ULP showed significant increase in accumulation of MTO1 gene product after treatment with anti-miR to miR-9-3p (Fig. 6B). The MTO1 mRNA is listed by TargetScanHuman as a target of miR-9-3p [31]. Significant changes in GTPBP3 or TRMU were not detected by Western blot after treatment of TOV-LT cells with anti-miRs-9-5p or anti-miR-9-3p. The GTPBP3 transcript is listed by TargetsScanHuman as a target of miR-9-5p; TRMU is not listed as a direct target of miR-9.

**Figure 6.**
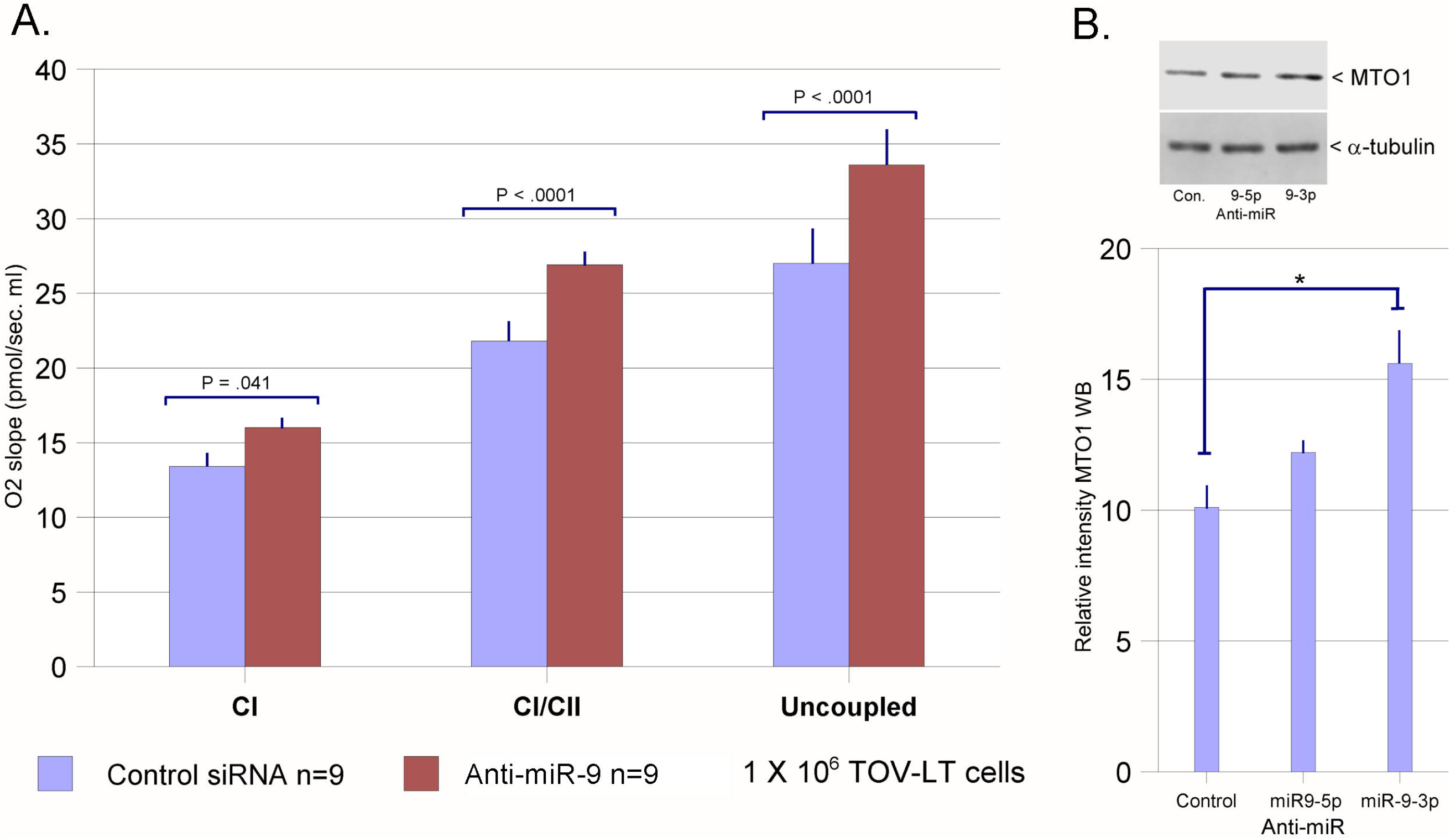
A. Mean values of nine independent respirometry determinations for TOV-LT treated with miR-9 (5p + 3p) or control siRNAs. B. Densitometry analyses of Western blots for MTO1 gene product in TOV-LT cells treated with miR-9 anti-miRs. An example of the blotted samples is shown, with mean values determined from three independent data points for control and each anti-miR produced on the same blot.

### Spheroid formation enhances intraperitoneal survival of TOV-112D ovarian carcinoma cells

To determine how spheroid formation affects the intraperitoneal survival of mobile tumor cells, 10^7^ TOV-LT or CAP cells growing in log phase over ULP were injected into the peritoneum of immunocompromised NOD/SKID mice. To recapitulate accurately the challenges of survival in the peritoneal microenvironment, the cells were injected in PBS lacking matrix materials (e.g., Matrigel). The cells had been modified to express luciferase, and survival was assessed every five days by in vivo imaging. The mean signal intensity in six animals injected with either cell type was equivalent five days after injection, but at 10, 15 and 20 days CAP cells showed significantly enhanced survival (Fig. 7). At 30-40 days signal could no longer be detected. The data suggest that the increased OXPHOS activity afforded by the spheroid architecture of CAP cells enhanced their survival compared to TOV-LT cells in the nutrient poor environment of the peritoneum.

**Figure 7.**
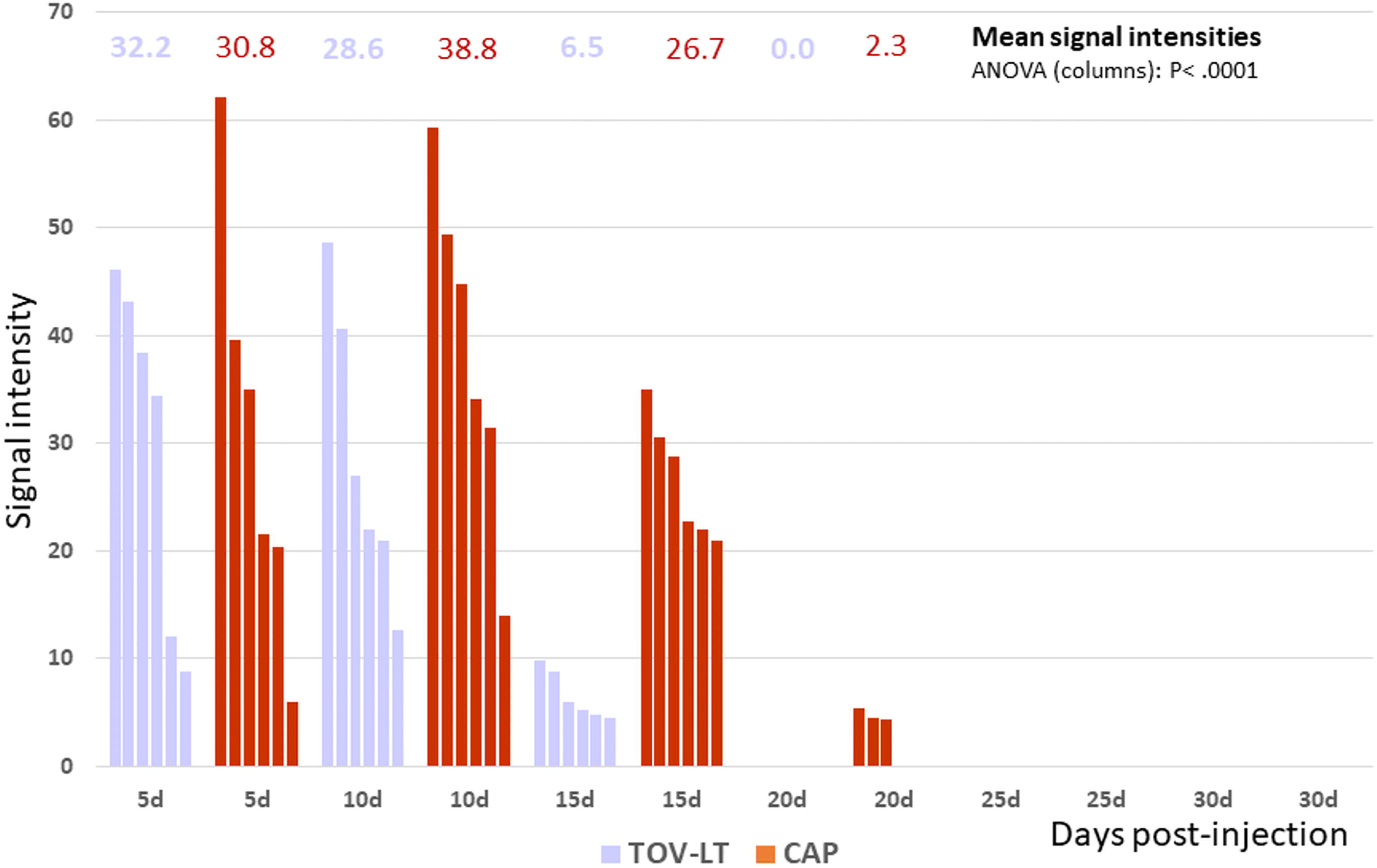
Comparison of intraperitoneal survival rates of TOV-LT and CAP cells. TOV-LT and CAP cells cultured over ULP were injected in logarithmic growth phase into NOD/SKID mice. Ten million cells were injected per mouse. The cells had been modified to express luciferase and were imaged in vivo every five days for the course of the experiment. Overall photo emissions per second for each animal are reported, and ANOVA was performed on mean emission values for the course of the experiment.

## Discussion

### Spheroid architecture promotes OXPHOS

The present studies aim to directly address the question of how the formation of spheroids impacts mitochondria function. Previous studies using mitochondrial inhibitors have indicated that mitochondrial activity is essential for spheroid formation, and that intracellular ATP depletion leads to loss of spheroidal architecture [32]. Considering the importance of ATP and GTP levels in controlling the assembly of cytoskeletal elements, this observation is not surprising. A previous report has associated a high bioenergetics profile with the ability of different ovarian cancer cell lines to form spheroids in culture [17]. That study surveyed several distinct cell lines, and concluded that cells showing a relatively elevated bioenergetics profile display enhanced ability to form spheroids. While this observation is consistent with and even supportive of our results, the suggestion that elevated bioenergetics drives spheroid formation diverges from the conclusions of our studies, which indicate instead that it is the formation of spheroids that enhances the OXPHOS activity of constituent cells. Of course, these conclusions are not mutually exclusive, and both mechanisms are likely relevant to tumor growth and metastases under specific conditions. The results of our studies are also consistent with the previous observation that 25 mitochondrial-related proteins show increased expression in 3D-mammospheres [32].

In the present studies, respirometry was performed on derivatives of TOV-112D cells that stably form (CAP) or do not form (TOV-LT) spheroids when grown over ULP. This approach allowed comparison of cells derived from the same tumor growing at the same rate under the same conditions (over ULP), except one set of cells was growing as spheroids (CAP) and the other was growing as dispersible aggregates (TOV-LT). The derivatives of TOV-112D reflect equivalent states exhibited by early passage TOV-112D cells when transitioned from TCP to ULP. Early passage TOV-112D cells growing over ULP (TOV-1) show frequent spheroid formation, but spheroid formation is lost after five passages over ULP (TOV-LT). We found that a clone of TOV-112D cells derived in a previous study (CAP cells; [25] stably forms spheroids for several generations over ULP. High-resolution respirometry with readily performed with the Oxygraph-2k system as the cells were growing in suspension. Analyses showed greater state 3 Complex I, I/II and uncoupled (state 3U) respiration rates per cell in CAP cell spheroids than in loose TOV-LT cells aggregates. Further studies showed that interference with spheroid architecture by treatment of CAP cells with N-cadherin siRNA reduced OXPHOS activity. Together, these results indicate that growth in suspension as spheroids promotes OXPHOS activity in comparison to growth as loose aggregates of cells. An essential connection between cytoskeletal architecture and mitochondrial function in cancer cells has been observed in previous studies [33]. Spheroid architecture likely impacts the underlying cellular cytoskeletal organization, resulting in positive regulation of mitochondrial functions and/or expression of mitochondrial proteins. Because of the activating defect in *CTNNB1* in TOV-112D cells, differences in gene expression or metabolism observed between spheroidal and non-spheroidal TOV-112D cells are more likely to be associated with cellular architecture than with alterations of β-catenin signaling due to differences in N-cadherin ligation [34, 35]. The effects of spheroid architecture on cytoskeletal and other intracellular structures may directly impact mitochondrial function. Spheroid architecture also may mediate changes in gene expression that impact metabolism and other aspects of cell growth and phenotype via a mechanism that alters the three-dimensional organization of chromatin in the nucleus [36].

### Differential expression of miR-221/222 and miR-9 in spheroids and non-spheroidal cells growing in suspension

Deep sequencing of small RNA libraries prepared from TOV-LT and CAP cells grown over ULP revealed potentially differences in expression of miRs. MicroRNAs showing large differences were further tested by qRT-PCR for significance and magnitude of difference. Expression of miR-221/222 was found in the initial screen and by qRT-PCR to be profoundly enhanced during growth as spheroids (CAP) as compared to loose aggregates (TOV-LT) over ULP. The miR-221/222 cluster encodes oncomirs that target cell cycle inhibitors and promote entry into S-phase [37]. In ovarian carcinomas, expression of miR-221/222 is positively associated with poor therapeutic outcomes [38], resistance to chemotherapeutics including cisplatin [39, 40], and maintenance of the cancer stem cell phenotypes [41–43]. Both drug-resistance and growth of cancer stem cells have been linked with tumor cell spheroid formation [44], and upregulation of miR-221/222 may be part of the mechanism by which these properties are conferred to tumor cells that comprise spheroids. Down-regulation of miR-9 was observed in CAP cells growing over ULP as spheroids compared to TOV-LT cells growing as loosely aggregated clumps. Published targets of miR-9 include mitochondrial t-RNA modifying enzymes (GTPBP3, MTO1, TRMU), and increased expression of miR-9 is involved in mitochondrial pathogenesis in MELAS (mitochondrial encephalopathy, lactic acidosis, and stroke-like episodes) syndrome [30]. We found that lipofection of anti-miR-9 (5p + 3p) into TOV-LT cells increased OXPHOS activity, consistent with reduced expression of miR-9 as a component of the mechanism of enhancing OXPHOS activity in spheroids. Western blot analyses of TOV-LT cells showed increased accumulation of MTO1 after treatment with miR9-3p antimiR, consistent with the analysis of miR9-3p. Changes in GTPBP3 or TRMU were not observed using either antimir, and while GTPBP3 is a predicted target of miR-9-5p, other studies have suggested that mir-9-5p expression may be involved in metabolic reprogramming that supports mitochondrial activity [45].

### Survival advantages of spheroid formation for mobile intraperitoneal ovarian carcinoma cells

In a glucose-deprived environment such as the peritoneal cavity, ovarian cancer stem cells adapt to maintain an oxidative phenotype. Isolated as effusions from epithelial ovarian cancer patients, ex vivo populations of ovarian cancer cells expressing stem cell markers display a metabolic profile characterized by preferential use of glucose through OXPHOS [46]. Under these conditions ovarian cancer stem cells show reliance on OXPHOS and enrichment of electron transport chain components [47]. Adaptation to glucose-deprived condition of the peritoneum by switching to oxidative metabolism is thought to maximize the survival of ovarian cancer stem cells and promote metastasis and tumor growth at distal sites. Consistent with this hypothesis, segregation of ovarian cancer stem cells with higher relative mitochondrial transmembrane potential yields cells with more potent tumor-initiating and spheroid-forming properties [48]. Using an immunodeficient murine xenograft model, we directly compared the survival of CAP cells injected into the peritoneum after growth over ULP as spheroids with the survival of TOV-LT cells injected into the peritoneum after growth over ULP as loose clumps. To better model the actual environmental conditions encountered by mobile (metastasizing) ovarian cancer cells, these experiments were performed in the absence of added matrix materials. The average duration of intraperitoneal survival of CAP cells exceeded that of TOV-LT by more than 5 days, consistent with the greater oxidative metabolism of CAP spheroids improving viability in the glucose-deprived environment of the peritoneum.

A limitation of these experiments was the inability of injected CAP or TOV-LT cultures to produce tumors or intraperitoneal carcinomatosis in the 40-day time period during which the experiment was conducted. The inability of a single dose of cultured ovarian cancers cells in the absence of supporting matrix to form intraperitoneal tumors in mice is consistent with the observations that development of cell lines that aggressively form peritoneal tumors or carcinomatosis often requires repeated passage through animals [49, 50]. Other studies have shown only moderate growth of TOV-112D as tumors in nude mice [51] and long-term culturing over ULP has probably reduced the tumorigenicity of CAP and TOV-LT cells. A caveat to enhancing tumorigenicity of TOV-LT and CAP cell by passaging through immunodeficient mice is that the process will also likely alter the spheroidogenic and metabolic properties of the cells as well. While the experiments presented herein provide direct evidence of enhanced survival of mobile tumor cells, future studies are in progress to examine the properties of tumorigenic cell lines that have been altered to express different oxidative and spheroidogenic properties.

Resolving how enhancement of OXPHOS activity by tumor spheroid formation influences cancer stem cell formation is a complex issue. Pluripotent stem cells (normal) and bulk tumor cells generally show glycolytic phenotype that is related both the generation of precursors for anabolic metabolism and the demands of rapidly growing cells [52–55]. Pluripotent stem cells switch to oxidative metabolism upon differentiation to multipotent, unipotent or terminal forms. For bulk tumor cells the persistence of glycolytic metabolism under aerobic conditions has been referred to as the Warburg effect or anaerobic glycolysis, and is the prototypical metabolic alteration that is considered to be a core hallmarks of cancer [56]. There is growing evidence that metastasized ovarian tumors metabolized fatty acids for energy production, and that ovarian cancer stem cells show enhanced capacity for OXPHOS in comparison to bulk tumors cells [57]. Studies of cancer stem cells have found that metabolic phenotypes vary between OXPHOS and glycolytic modes depending on the extracellular environment and the specific oncogenes and/or tumor suppressors that are driving tumorigenesis [58, 59]. Consistent with this, adaptation to a hypoxic environment, such as the milieu within a dense solid tumor, promotes a glycolytic phenotype among cancer stem cells [60, 61]. Understanding the mechanisms by which spheroid architecture affects ovarian cancer cell metabolic properties and gene expression will provide a basis for development of next generation anti-cancer therapeutics to impede metastasis and improve the efficacy of inductive and other chemotherapeutics.

## Conclusions

The results of these studies show that growth of an epithelial ovarian cancer cell line as spheroids induces greatly enhanced expression of miR-221/222, reduces expression of miR-9 and enhances OXPHOS activity. Growth as spheroids and the associated increase in OXPHOS activity promotes survival of the tumor cells in the nutrient-poor environment of the peritoneum, suggesting a new mechanism by which spheroid formation can enhance the metastatic potential of mobile tumor cells.

## Availability of data and materials

All data analyzed during this study are included in this published article and its supplementary information files. Associated raw data are available from the corresponding author on reasonable request.

### Abbreviations

*Adh*: adherent
*CSC*: Cancer stem cells
*DMEM*: Dulbecco’s Modified Eagle’s medium
*Fig.*: Figure
*Figs.*: Figures
*JG-B*: Janus green B
*PBS*: Phosphate-buffered saline, PH 7.4
*miR*: MicroRNA
*N-cad*: N-cadherin
*qRT-PCR*: Quantitative reverse transcriptase polymerase chain reaction
OXPHOS: Oxidative phosphorylation
*Pyrmalate*: Pyruvate + malate
*RPMI*: Rosewell Park Memorial Institute 1640 medium
*Sus*: suspension
*TCP*: Tissue culture polystyrene
*ULP*: Ultralow attachment polystyrene

## Acknowledgements

The authors would like to thank Edward E. McKee and Itzel R. Gutierrez for material assistance, protocols, advise, and encouragement in the completion of these studies.

## Author information

### Contributions

ASW and DSK conceived and designed the experiments. The investigations, data collection, and data analysis were conducted by ASW, CNH, MOT and DSK. DSK wrote the manuscript and supervised the study.

Corresponding author: D. Stave Kohtz

## Ethics declarations

Ethics approval and consent to participate:

Not applicable.

Consent for publication:

Not applicable.

Competing interests:

The authors declare no competing interests.

## Supplementary Files

Supplementary Fig. 1. Comparison of OXPHOS activity in TOV-LT and CAP cells grown over ULP or after trypsinization from TCP.

Supplementary Fig. 2. Normalized counts for comparative HiSeq × 10 sequencing of an smRNA libraries prepared from TOV-LT and CAP cells.

