## Supplementary figures and images for "Spheroid architecture strongly induces miR-221/222 expression and promotes oxidative phosphorylation and survival of mobile tumor cells through a mechanism that includes restriction of miR-9 expression"

### Supplemental Figure 1

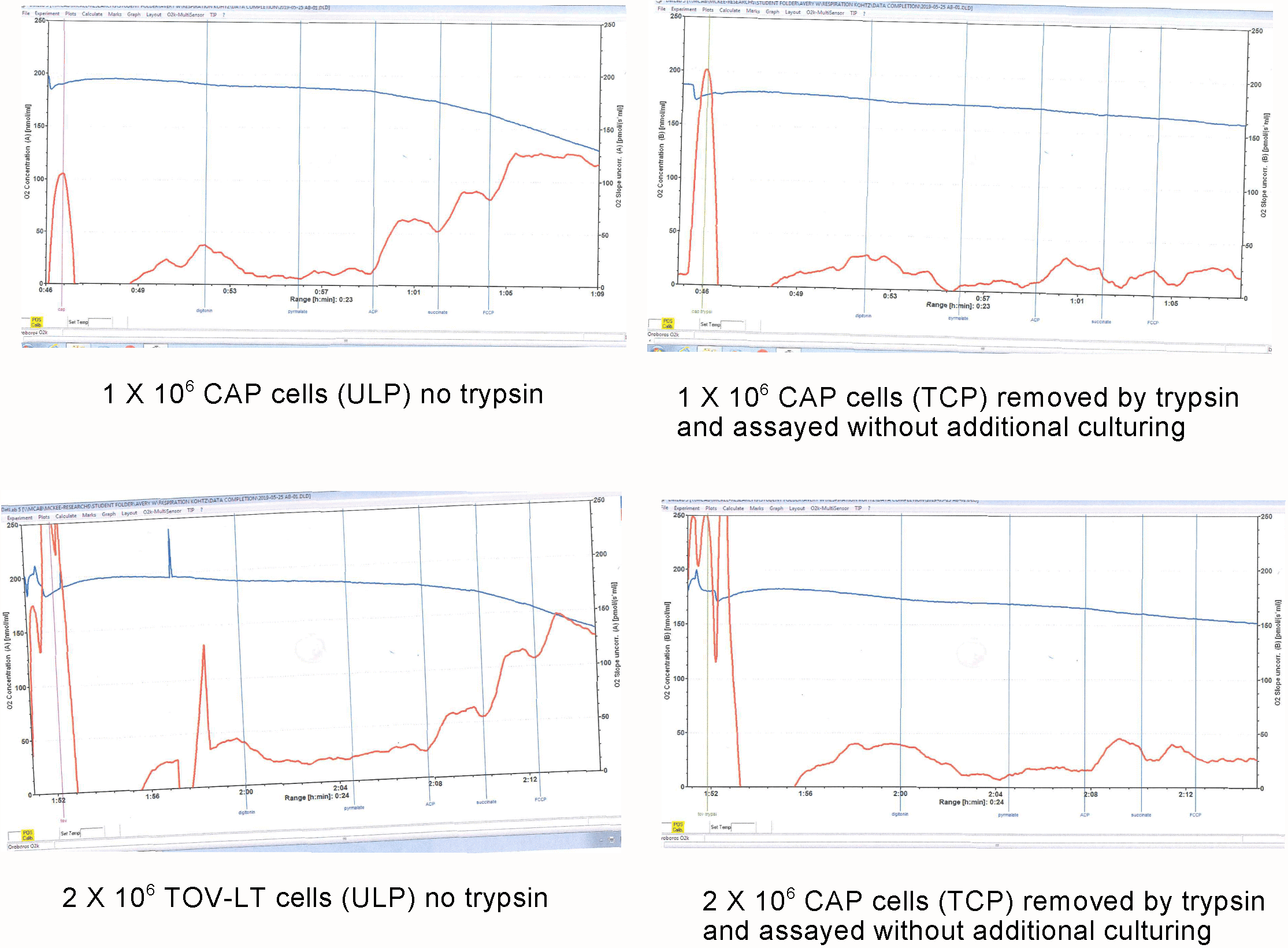
