## Supplemental Figure 2 for "Spheroid architecture strongly induces miR-221/222 expression and promotes oxidative phosphorylation and survival of mobile tumor cells through a mechanism that includes restriction of miR-9 expression"

| | CAP-se51 | TOV-LT-se51 |
|---|---|---|
| SampleGro | CAP | TOV-LT |
| let-7 | 702003.1 | 347871.4 |
| mir-1 | 5.92847 | 40.48262 |
| mir-10 | 221926.3 | 111264 |
| mir-101 | 617.7465 | 753.9889 |
| mir-103 | 61239.9 | 98569.28 |
| mir-105 | 3.557082 | 19.39792 |
| mir-1179 | 0 | 12.65082 |
| mir-1180 | 9374.096 | 3861.03 |
| mir-122 | 15.41402 | 0 |
| mir-1224 | 7.114163 | 0 |
| mir-1225 | 20.1568 | 2.530164 |
| mir-1226 | 177.8541 | 83.49541 |
| mir-1228 | 0 | 5.060328 |
| mir-124 | 2.371388 | 125.6648 |
| mir-1243 | 0 | 6.747104 |
| mir-1245 | 28.45665 | 59.88055 |
| mir-1246 | 40.31359 | 454.5861 |
| mir-1247 | 1.185694 | 0 |
| mir-1248 | 4.742776 | 9.277268 |
| mir-1249 | 322.5087 | 323.861 |
| mir-1250 | 64.02747 | 115.5442 |
| mir-1254 | 94.85551 | 47.22973 |
| mir-1255 | 3.557082 | 1.686776 |
| mir-1257 | 2.371388 | 15.18098 |
| mir-126 | 419.7356 | 1486.05 |
| mir-1260a | 36.75651 | 8.43388 |
| mir-1260b | 5476.72 | 7060.844 |
| mir-1261 | 0 | 0.843388 |
| mir-1262 | 4.742776 | 32.89213 |
| mir-1266 | 0 | 3.373552 |
| mir-1267 | 0 | 14.3376 |
| mir-1269 | 0 | 2.530164 |
| mir-127 | 248.9957 | 546.5154 |
| mir-1270 | 4.742776 | 3.373552 |
| mir-1271 | 666.36 | 585.3113 |
| mir-1272 | 9.485551 | 58.19377 |
| mir-1273 | 29.64235 | 37.10907 |
| mir-1273c | 5.92847 | 24.45825 |
| mir-1275 | 3.557082 | 10.12066 |
| mir-1276 | 47.42776 | 43.85618 |
| mir-1277 | 1.185694 | 0 |
| mir-1278 | 0 | 6.747104 |
| mir-128 | 7019.308 | 5035.026 |
| mir-1285 | 16.59971 | 53.97683 |
| mir-1286 | 0 | 15.18098 |

| | | |
|---|---|---|
| mir-1287 | 4.742776 | 0 |
| mir-1288 | 17.78541 | 9.277268 |
| mir-1289 | 3.557082 | 4.21694 |
| mir-129 | 295.2378 | 164.4607 |
| mir-1291 | 17.78541 | 17.71115 |
| mir-1292 | 118.5694 | 106.2669 |
| mir-1294 | 5.92847 | 10.96404 |
| mir-1295 | 49.79914 | 14.3376 |
| mir-1296 | 895.1989 | 543.1419 |
| mir-1298 | 199.1966 | 18.55454 |
| mir-1299 | 0 | 1.686776 |
| mir-130 | 4696.534 | 4372.967 |
| mir-1301 | 7468.686 | 5218.885 |
| mir-1302 | 9.485551 | 5.903716 |
| mir-1303 | 5092.555 | 2790.771 |
| mir-1304 | 104.3411 | 266.5106 |
| mir-1305 | 2.371388 | 0 |
| mir-1306 | 201.568 | 91.92929 |
| mir-1307 | 1686.057 | 1464.122 |
| mir-132 | 3676.837 | 16152.57 |
| mir-1322 | 0 | 14.3376 |
| mir-1323 | 0 | 9.277268 |
| mir-133 | 14.22833 | 41.32601 |
| mir-134 | 82.99857 | 331.4515 |
| mir-1343 | 26.08527 | 19.39792 |
| mir-135 | 0 | 13.49421 |
| mir-136 | 0 | 0.843388 |
| mir-137 | 7.114163 | 14.3376 |
| mir-138 | 11.85694 | 10.12066 |
| mir-139 | 69.95594 | 80.12186 |
| mir-140 | 3012.848 | 2326.907 |
| mir-142 | 522.891 | 367.7172 |
| mir-143 | 34.38512 | 19.39792 |
| mir-146 | 1032.739 | 756.519 |
| mir-1468 | 81.81288 | 101.2066 |
| mir-147 | 0 | 0.843388 |
| mir-148 | 3670.908 | 3597.893 |
| mir-149 | 3306.9 | 1070.259 |
| mir-15 | 41001.3 | 40260.81 |
| mir-150 | 2.371388 | 12.65082 |
| mir-1538 | 1.185694 | 0 |
| mir-154 | 1036.296 | 4310.556 |
| mir-155 | 297.6092 | 220.9677 |
| mir-17 | 53279.16 | 67940.81 |
| mir-181 | 173688.7 | 86946.55 |
| mir-182 | 3718.336 | 12334.55 |
| mir-183 | 11585.42 | 29457.01 |

| | | |
|---|---|---|
| mir-184 | 21.34249 | 16.02437 |
| mir-185 | 999.54 | 888.9309 |
| mir-186 | 5151.84 | 4270.917 |
| mir-187 | 8.299857 | 26.14503 |
| mir-188 | 6052.967 | 4652.128 |
| mir-19 | 237.1388 | 220.9677 |
| mir-190 | 15.41402 | 22.77148 |
| mir-1908 | 503.9199 | 178.7983 |
| mir-1909 | 2.371388 | 0 |
| mir-191 | 80589.24 | 39216.7 |
| mir-1910 | 177.8541 | 75.06153 |
| mir-1911 | 200.3823 | 6.747104 |
| mir-1912 | 11.85694 | 0 |
| mir-1913 | 0 | 4.21694 |
| mir-1914 | 5.92847 | 5.060328 |
| mir-1915 | 228.8389 | 27.8318 |
| mir-192 | 72.32733 | 149.2797 |
| mir-193 | 12011.08 | 2808.482 |
| mir-194 | 194.4538 | 186.3887 |
| mir-196 | 29903.2 | 18036.7 |
| mir-197 | 1913.71 | 844.2314 |
| mir-1976 | 0 | 0.843388 |
| mir-199 | 86090.86 | 117256.2 |
| mir-203 | 10.67125 | 28.67519 |
| mir-205 | 32.01374 | 0 |
| mir-21 | 19376.61 | 35277.23 |
| mir-210 | 305.909 | 437.7184 |
| mir-2110 | 392.4647 | 49.75989 |
| mir-2114 | 0 | 3.373552 |
| mir-214 | 153581.7 | 119285.4 |
| mir-216 | 0 | 8.43388 |
| mir-218 | 33.19943 | 165.304 |
| mir-219 | 47.42776 | 21.92809 |
| mir-22 | 3723.079 | 3166.922 |
| mir-221 | 28911.96 | 926.04 |
| mir-223 | 2.371388 | 4.21694 |
| mir-2278 | 15.41402 | 13.49421 |
| mir-23 | 103066.4 | 50869.79 |
| mir-2355 | 59.2847 | 40.48262 |
| mir-2392 | 48.61345 | 106.2669 |
| mir-23c | 26.08527 | 11.80743 |
| mir-24 | 41902.42 | 33703.47 |
| mir-25 | 338314 | 249297.9 |
| mir-26 | 95245.61 | 68833.95 |
| mir-27 | 26213.32 | 21334.34 |
| mir-28 | 26231.11 | 24236.44 |
| mir-29 | 29612.71 | 10939.59 |

| | | |
|---|---|---|
| mir-296 | 1230.75 | 570.1303 |
| mir-298 | 13.04263 | 0.843388 |
| mir-299 | 0 | 23.61486 |
| mir-30 | 71004.09 | 73282.83 |
| mir-3064 | 16.59971 | 18.55454 |
| mir-3065 | 14.22833 | 48.9165 |
| mir-3074 | 82.99857 | 83.49541 |
| mir-3121 | 16.59971 | 0.843388 |
| mir-3126 | 7.114163 | 36.26568 |
| mir-3127 | 68.77025 | 59.03716 |
| mir-3129 | 211.0535 | 174.5813 |
| mir-3130 | 22.52818 | 0 |
| mir-3135 | 40.31359 | 38.79585 |
| mir-3136 | 1.185694 | 0 |
| mir-3138 | 26.08527 | 0 |
| mir-3140 | 7.114163 | 24.45825 |
| mir-3145 | 0 | 0.843388 |
| mir-3146 | 9.485551 | 0 |
| mir-3150 | 0 | 13.49421 |
| mir-3154 | 0 | 5.060328 |
| mir-3155 | 0 | 0.843388 |
| mir-3158 | 36.75651 | 53.97683 |
| mir-3170 | 3.557082 | 5.903716 |
| mir-3173 | 193.2681 | 129.8818 |
| mir-3174 | 0 | 0.843388 |
| mir-3179 | 13.04263 | 12.65082 |
| mir-3180 | 8334.242 | 4208.506 |
| mir-3192 | 0 | 8.43388 |
| mir-3198 | 9.485551 | 0.843388 |
| mir-3199 | 27.27096 | 0 |
| mir-32 | 77.0701 | 139.159 |
| mir-320 | 26316.48 | 24989.59 |
| mir-3200 | 345.0369 | 492.5386 |
| mir-322 | 1891.182 | 1274.359 |
| mir-324 | 3848.762 | 2024.131 |
| mir-326 | 317.766 | 273.2577 |
| mir-328 | 2238.59 | 313.7403 |
| mir-329 | 333.18 | 1303.034 |
| mir-33 | 318.9517 | 368.5606 |
| mir-330 | 1005.468 | 2508.236 |
| mir-331 | 2972.535 | 3028.606 |
| mir-335 | 58.099 | 95.30284 |
| mir-337 | 0 | 17.71115 |
| mir-338 | 45.05637 | 164.4607 |
| mir-339 | 793.2292 | 904.9553 |
| mir-34 | 41.49929 | 94.45945 |
| mir-340 | 221.7248 | 912.5458 |

| | | |
|---|---|---|
| mir-342 | 2180.491 | 4551.765 |
| mir-345 | 1167.909 | 3904.886 |
| mir-3605 | 40.31359 | 62.41071 |
| mir-3607 | 1236.679 | 931.9437 |
| mir-361 | 7666.697 | 4468.27 |
| mir-3612 | 1.185694 | 169.521 |
| mir-3613 | 47.42776 | 34.57891 |
| mir-3615 | 234.7674 | 191.4491 |
| mir-362 | 2128.321 | 2641.491 |
| mir-363 | 0 | 133.2553 |
| mir-3648 | 0 | 2.530164 |
| mir-365 | 2283.646 | 1470.025 |
| mir-3661 | 8.299857 | 0 |
| mir-3664 | 1.185694 | 0 |
| mir-368 | 3.557082 | 82.65202 |
| mir-3680 | 0 | 7.590492 |
| mir-3688 | 0 | 5.903716 |
| mir-370 | 673.4741 | 1514.725 |
| mir-374 | 460.0492 | 661.2162 |
| mir-375 | 43.87067 | 336.5118 |
| mir-378 | 6751.341 | 12000.57 |
| mir-378_2 | 0 | 0.843388 |
| mir-379 | 139.9119 | 723.6269 |
| mir-383 | 36.75651 | 0 |
| mir-3912 | 0 | 0.843388 |
| mir-3934 | 5.92847 | 3.373552 |
| mir-3938 | 0 | 4.21694 |
| mir-3940 | 43.87067 | 3.373552 |
| mir-412 | 46.24206 | 330.6081 |
| mir-422 | 0 | 5.903716 |
| mir-423 | 158326.9 | 88984.18 |
| mir-425 | 9308.883 | 6244.445 |
| mir-431 | 7.114163 | 200.7263 |
| mir-432 | 35.57082 | 163.6173 |
| mir-433 | 77.0701 | 186.3887 |
| mir-4421 | 8.299857 | 53.97683 |
| mir-4429 | 0 | 1.686776 |
| mir-4444 | 7.114163 | 0 |
| mir-4449 | 0 | 5.903716 |
| mir-4477 | 1.185694 | 17.71115 |
| mir-448 | 39.1279 | 0 |
| mir-449 | 3.557082 | 4.21694 |
| mir-450 | 120.9408 | 242.0524 |
| mir-4510 | 97.2269 | 80.12186 |
| mir-4524 | 0 | 0.843388 |
| mir-454 | 313.0232 | 777.6037 |
| mir-455 | 424.4784 | 1092.187 |

| | | |
|---|---|---|
| mir-4637 | 1.185694 | 0 |
| mir-4654 | 0 | 6.747104 |
| mir-4659 | 4.742776 | 19.39792 |
| mir-4660 | 1.185694 | 7.590492 |
| mir-4662 | 0 | 12.65082 |
| mir-4677 | 3.557082 | 0 |
| mir-4738 | 0 | 0.843388 |
| mir-4742 | 8.299857 | 6.747104 |
| mir-4743 | 1.185694 | 0 |
| mir-4766 | 0 | 1.686776 |
| mir-4791 | 3.557082 | 6.747104 |
| mir-483 | 0 | 11.80743 |
| mir-484 | 4526.979 | 3414.878 |
| mir-485 | 150.5831 | 375.3077 |
| mir-486 | 26.08527 | 36.26568 |
| mir-491 | 194.4538 | 177.9549 |
| mir-493 | 125.6836 | 699.1686 |
| mir-497 | 245.4386 | 192.2925 |
| mir-499 | 0 | 5.060328 |
| mir-500 | 8323.571 | 9047.866 |
| mir-503 | 1001.911 | 1472.555 |
| mir-504 | 91.29843 | 13.49421 |
| mir-505 | 5933.212 | 3184.633 |
| mir-506 | 37.94221 | 51.44667 |
| mir-515 | 27.27096 | 49.75989 |
| mir-541 | 0 | 3.373552 |
| mir-542 | 18.9711 | 23.61486 |
| mir-544 | 0 | 2.530164 |
| mir-548 | 113.8266 | 199.0396 |
| mir-549 | 0 | 2.530164 |
| mir-550 | 77.0701 | 176.2681 |
| mir-551 | 2934.592 | 818.9297 |
| mir-5583 | 0 | 3.373552 |
| mir-561 | 0 | 0.843388 |
| mir-572 | 4.742776 | 0 |
| mir-573 | 1.185694 | 3.373552 |
| mir-574 | 1214.151 | 293.499 |
| mir-576 | 15.41402 | 44.69956 |
| mir-577 | 1.185694 | 2.530164 |
| mir-582 | 75.88441 | 81.80863 |
| mir-588 | 0 | 2.530164 |
| mir-589 | 883.342 | 759.0492 |
| mir-590 | 24.89957 | 29.51858 |
| mir-597 | 14.22833 | 16.86776 |
| mir-598 | 104.3411 | 222.6544 |
| mir-599 | 6266.392 | 34281.19 |
| mir-600 | 1.185694 | 0 |

| | | |
|---|---|---|
| mir-605 | 2.371388 | 0 |
| mir-6134 | 1.185694 | 0 |
| mir-615 | 17117.86 | 10262.35 |
| mir-616 | 29.64235 | 57.35038 |
| mir-624 | 17.78541 | 16.86776 |
| mir-625 | 79.44149 | 147.5929 |
| mir-627 | 0 | 2.530164 |
| mir-628 | 139.9119 | 199.0396 |
| mir-629 | 151.7688 | 1248.214 |
| mir-632 | 0 | 1.686776 |
| mir-636 | 1.185694 | 5.903716 |
| mir-639 | 5.92847 | 0 |
| mir-641 | 37.94221 | 78.43508 |
| mir-642 | 67.58455 | 14.3376 |
| mir-6505 | 11.85694 | 15.18098 |
| mir-651 | 8.299857 | 4.21694 |
| mir-6511 | 349.7797 | 242.8957 |
| mir-6516 | 42.68498 | 29.51858 |
| mir-652 | 14440.57 | 7766.76 |
| mir-654 | 67.58455 | 256.3899 |
| mir-657 | 0 | 4.21694 |
| mir-664 | 36.75651 | 22.77148 |
| mir-665 | 7.114163 | 16.02437 |
| mir-668 | 0 | 4.21694 |
| mir-671 | 413.8072 | 237.8354 |
| mir-676 | 4.742776 | 0.843388 |
| mir-6770 | 1.185694 | 0.843388 |
| mir-6794 | 9.485551 | 9.277268 |
| mir-6859 | 1.185694 | 0 |
| mir-6862 | 1.185694 | 13.49421 |
| mir-7 | 1057.639 | 3149.211 |
| mir-744 | 6733.556 | 3088.487 |
| mir-760 | 347.4083 | 549.889 |
| mir-766 | 193.2681 | 117.2309 |
| mir-767 | 10.67125 | 17.71115 |
| mir-769 | 1970.623 | 1563.641 |
| mir-770 | 0 | 7.590492 |
| mir-8 | 382.9791 | 333.9816 |
| mir-873 | 687.7025 | 141.6892 |
| mir-874 | 32.01374 | 7.590492 |
| mir-875 | 186.1539 | 777.6037 |
| mir-876 | 22.52818 | 9.277268 |
| mir-877 | 685.3311 | 1424.482 |
| mir-885 | 3.557082 | 0 |
| mir-887 | 244.2529 | 166.9908 |
| mir-889 | 0 | 39.63924 |
| mir-891 | 9.485551 | 0 |

| | | |
|---|---|---|
| mir-9 | 801.5291 | 12365.75 |
| mir-935 | 113.8266 | 121.4479 |
| mir-937 | 34.38512 | 56.507 |
| mir-939 | 8.299857 | 1.686776 |
| mir-940 | 718.5305 | 463.02 |
| mir-941 | 282.1952 | 200.7263 |
| mir-942 | 59.2847 | 38.79585 |
| mir-943 | 0 | 3.373552 |
| mir-944 | 4.742776 | 5.060328 |
| mir-95 | 1938.61 | 1518.942 |
| mir-96 | 193.2681 | 313.7403 |
